# Multi-spectral Fluorescent Reporter Influenza A Viruses Allow for *in vivo* Studies of Innate Immune Function in Zebrafish

**DOI:** 10.1101/2023.10.31.564888

**Authors:** Brandy-Lee Soos, Alec Ballinger, Mykayla Weinstein, Haley Foreman, Julianna Grampone, Samuel Weafer, Connor Aylesworth, Benjamin L. King

## Abstract

Influenza virus infection can cause severe respiratory disease and is estimated to cause millions of illnesses annually. Studies of the contribution of the innate immune response to influenza A virus (IAV) to viral pathogenesis may yield new antiviral strategies. Zebrafish larvae are useful models to study the innate immune response to pathogens, including IAV, *in vivo*. Here, we demonstrate how Color-flu, four fluorescent IAV strains originally developed for mice, can be used to study host-virus interactions by simultaneously monitoring virus particles, neutrophils, and macrophages *in vivo*. Using this model, we show how the angiotensin-converting enzyme inhibitor, ramipril, and mitophagy inhibitor, MDIVI-1, improved survival, decreased viral burden, and improved the respiratory burst response to IAV infection. The Color-flu zebrafish model of IAV infection is complementary to other models as it is the only model where interactions between virus particles and host cells in an intact vertebrate can be visualized *in vivo*.

## INTRODUCTION

Influenza A virus (IAV) infection can result in acute respiratory inflammation that requires hospital care, and in severe cases can lead to death. An estimated 10-37 million influenza infections occur each year in the US, with 114,000-624,000 needing hospital care for associated respiratory and heart symptoms[1]. 5,000-27,000 deaths occur each year in the US from influenza infections[2]. While vaccines have been widely used, they have only been between 19-60% effective in preventing influenza because of antigenic variation in strains circulating in different populations and the ability to formulate vaccines against those strains[3, 4]. Outbreaks of new influenza strains can become pandemic, such as the 2009 A(H1N1)pdm09 strain that resulted in an estimated 60.8 million infections, 274,000 hospitalizations, and 12,000 deaths in the US[5]. Antiviral therapies, such as oseltamivir (Tamiflu®), are valuable tools to treat influenza but the potential risk that drug-resistant strains pose make it necessary to develop new therapies that target alternative mechanisms.

One roadblock to understanding the pathogenesis of influenza infection is the relative contributions of the virus and host factors have not been well characterized *in vivo*. Mammalian influenza virus infections originate in respiratory epithelial cells and alveolar macrophages[6]. The innate immune response to influenza infection includes type I interferons (IFN) and proinflammatory cytokines and chemokines that activate neutrophils and macrophages. The innate antiviral immune response is activated in response to influenza infection and initiates a feed-forward loop that leads to the recruitment of excess neutrophils characterized by dysregulation of IFN expression and reactive oxidative species (ROS) production[7]. Characterizing the complex dynamics of host immune cells during the response to influenza virus infection requires simultaneous monitoring of multiple cell types along with the virus. Biopsy studies in human, primate or mammalian models cannot show the temporal dynamics of viral invasion and subsequent neutrophil and macrophage recruitment.

Influenza viruses that express fluorescent proteins are powerful tools to understand viral pathogenesis and host-virus interactions *in vivo*. A green fluorescent protein (GFP) reporter strain of A/Puerto Rico/8/34 (PR8; H1N1) was used to study antigen presentation during IAV infection[8]. Fukuyama *et al*. generated four different mouse-adapted PR8 strains that express different fluorescent reporter proteins[9]. These “Color-flu” strains express either Venus (mVenus-PR8), a GFP with improved chromophore formation and brightness, enhanced cyan fluorescent protein (eCFP-PR8), enhanced GFP (eGFP-PR8), or mCherry (mCherry-PR8). Importantly, the development of these mouse-adapted strains included serial passaging and selecting for strains that had increased pathogenicity and strong fluorescent protein expression to overcome the attenuation observed with viruses expressing reporter proteins[10, 11].

The zebrafish is a powerful model to study host-virus interactions as genetic tools can be combined with *in vivo* imaging of transparent embryos[12]. Major human immune signaling pathways that respond to viral infection are conserved in zebrafish[12]. Gabor *et al*. established the zebrafish model of IAV infection[13]. In that study, it was demonstrated that: 1) zebrafish embryos express α-2,6-linked sialic acid receptors; 2) embryos have reduced survival following IAV disseminated infection; 3) IAV replicates in embryos; 4) interferon phi 1 (*ifnphi1*) and myxovirus (influenza) resistance A (*mxa*) had upregulated expression with IAV infection; 5) disseminated IAV infection resulted in necrosis of liver, gill and head kidney tissue along with pericardial edema; and 6) that pathological phenotypes from IAV infection were reduced in embryos treated with the neuraminidase inhibitor, Zanamivir. This study also showed how neutrophils are recruited to the site of localized infection in the swimbladder using fluorescent confocal imaging of the zebrafish embryos infected with NS1-GFP PR8 IAV[8]. These initial studies demonstrated that *in vivo* imaging of fluorescent influenza strains in a vertebrate is possible.

In this study, we demonstrate the application of Color-Flu in zebrafish embryos to study host-virus interactions *in vivo*. We first compared survival and viral burden of embryos to systemic infection by H1N1 PR8 with Color-Flu. Next, we show how microinjection of Color-Flu into the circulatory system results in disseminated infection throughout the embryo. We also show how Color-Flu can be simultaneously imaged with neutrophil and macrophage reporter lines. As genetic background can influence phenotypes, we examined survival of two wild-type zebrafish strains, AB and Ekkwill (EK), along with the pigment mutant, *casper* (*mitfa^w2/w2^; mpv17^a9/a9^*)[14, 15], with disseminated infection. Next, we demonstrate how ColorFlu can be used to test for the efficacy of two small molecules, ramipril and mitochondrial division inhibitor 1 (MDIVI-1), that we found to increase survival following systemic infection. Ramipril is an inhibitor of angiotensin-converting enzyme (ACE)[16], and a recent study found that individuals who have prescriptions for ACE inhibitors had a lower risk of influenza[17]. MDIVI-1 inhibits dynamin 1 like (DNM1L) and blocks apoptosis by preventing mitochondrial and peroxisomal division[18]. Together these studies demonstrate the utility of using Color-Flu in a zebrafish model of IAV infection to study host-virus interactions and screen small molecules *in vivo*.

## MATERIALS AND METHODS

### Zebrafish Care and Maintenance

Zebrafish used in this study were housed and maintained in the Zebrafish Facility at the University of Maine in accordance with the recommendations and standards in the Guide for the Care and Use of Laboratory Animals of the National Institutes of Health and the Institutional Animal Care and Use Committee (IACUC) at the University of Maine. Protocols utilized in this study were approved by the IACUC at the University of Maine (Protocol Number: A2021-02-02). Zebrafish were housed in recirculating tanks following standard procedures of a 14-hour light, 10-hour dark cycle at 28°C[19]. Zebrafish lines used in this study were AB, Ekkwill (EK), *casper* (*mitfa^w2/w2^; mpv17^a9/a9^*)[14], and Tg(*mpeg1:*eGFP*;lyz:*dsRed). Embryos were obtained by spawning adult zebrafish using varying sets of females. Embryos were kept at 33°C in 50 mL of sterilized egg water (60 μg/mL Instant Ocean Sea Salts; Aquarium Systems, Mentor, OH) in 100 mm × 25 mm petri dishes (catalog number 89220-696, VWR, Radnor, PA) with water changes every 2 days.

### MDCK/London Cell Culture

Madin-Dardy canine kidney/London (MDCK/London; Influenza Reagent Resource) cells (passage 3-4) were cultured using a modified protocol originally developed by Eisfield et al.[20]. Cells were grown at 37°C with 5% CO_2_ in T-175 flasks (CELLSTAR Flasks, USA Scientific, Ocala, FL) in minimal essential medium (MEM; Gibco catalog number 11090073, Thermo Fisher Scientific, Waltham, MA), containing final percentages/concentrations as follows: 5% heat-inactivated newborn calf serum (NCS; Gibco catalog number 26010074), 2% heat-inactivated fetal bovine serum (FBS; Gibco catalog number 16140071), 0.23% sodium bicarbonate solution (Gibco catalog number 25080094), 2% MEM amino acids (from 50X stock; Gibco catalog number 11130051), 1% MEM vitamin solution (from 100X stock; Gibco catalog number 11120052), 4 mM L-glutamine (Gibco catalog number 25030081), and 1% Antibiotic-Antimycotic (from 100X stock; Gibco catalog number 15240062). Cells were maintained by washing twice with 1× Dulbecco’s phosphate-buffered saline (PBS, pH 7.4), trypsinizing with 0.25% trypsin-EDTA with phenol red (Gibco catalog number 25300054), and passaged in a 1:10 dilution every 2-3 days up to passage eight. Virus-infected cells were grown in MEM-BSA-TPCK media[20] that was prepared similar to the MEM media described above but supplemented with 1 µg/mL Tosyl phenylalanyl chloromethyl ketone (TPCK) trypsin (Worthington Chemical Corporation, Lakewood, NJ) and with Bovine Albumin Fraction V (7.5% solution; Gibco catalog number 15260037) instead of NCS and FBS.

### Influenza Virus

PR8 influenza virus (A/PR/8/34 (H1N1) Purified Antigen; catalog number 10100374) was purchased from Charles River Laboratories (now AVS Bio in Norwich, CT). Upon arrival, the virus was defrosted on ice, aliquoted into microcentrifuge tubes, and stored at −80°. Prior to use, virus aliquots were thawed on ice and diluted in cold sterile Hank’s Buffered Salt Solution (HBSS) using a ratio of 87% virus to 13% diluent.

Color-flu[9] influenza virus strains MA-mVenus-PR8 (mVenus-PR8), MA-eCFP-PR8 (eCFP-PR8), MA-eGFP-PR8 (eGFP-PR8), and MA-mCherry-PR8 (mCherry-PR8) were kindly provided by Dr. Yoshihiro Kawaoka’s laboratory and stored at −80°C. Color-flu virus strains were propagated in separated T-25 flasks (CELLSTAR Flasks, USA Scientific) using MDCK/London cells using MEM-BSA-TPCK as outlined in Eisfield et al[20] (see MDCK/London Cell Culture section). Color-flu strains were grown for 4 days before being collected, filter sterilized using 0.45 µm tube top vacuum filters (VWR), aliquoted, and stored at −80°C until use.

### Microinjection

Microinjection was used to inject either vehicle (HBSS) controls or influenza virus to introduce a disseminated or localized infection in zebrafish larvae. First, influenza virus strains were thawed on ice for 30 minutes. Virus strains were diluted in cold, sterile HBSS (Gibco catalog number 14170120) at 87% virus and 13% HBSS. Next, larvae were anesthetized in sodium bicarbonate-buffered MS-222 (Tricaine methanesulfonate) (Syndel, Ferndale, WA) solution (200 mg/L) at 2 or 3 dpf for disseminated infection experiments, or 4 dpf for localized infection experiments. Larvae were then lined up on a 2% agarose gel in a Petri dish coated with 3% methylcellulose. Microinjections of virus or HBSS were conducted using pulled microcapillary pipettes (Pulled microcapillary pipettes (1.2 mm outside diameter, 0.94 mm inside diameter, Sutter Instruments,

Novato, CA) controlled with an MPPI-3 pressure microinjector (Applied Scientific Instruments, Eugene, OR). For disseminated infection experiments of PR8 virus, 1 or 2 nL of virus or HBSS was injected into the duct of Cuvier (DC) at 2 or 3 dpf, respectively. 6 nL of Color-flu virus or HBSS was injected into the DC at 3 dpf for disseminated Color-flu infection experiments. For localized infection experiments with PR8 virus, 4 nL of virus or HBSS was injected into the swimbladder of larvae at 4 dpf. 10 nL of Color-flu or HBSS was injected into the swimbladder at 4 dpf for localized Color-flu infection experiments. For control experiments, larvae were also injected with heat-inactivated or UV-inactivated virus. Virus aliquots were either heat-inactivated at 80°C for 5 minutes, or UV-inactivated with 254 nm light exposure for 60 minutes on ice (UV CrossLinker, VWR). For virus infections, microcapillary pipettes were changed hourly to keep the virus viable. For disseminated infection experiments, zebrafish were sorted into petri dishes at a density of ∼50 larvae/dish and maintained in embryo water at 33°C (see Zebrafish section). For localized infection experiments, zebrafish were kept at a density of ∼45 larvae/dish.

### Drug Exposures

For all small molecule drug studies, virus-infected or HBSS-injected larvae were exposed to either dimethyl sulfoxide (DMSO; Sigma-Aldrich, St. Louis, MO), DMSO-solubilized ramipril (Cayman Chemical Company, Ann Arbor, MI), or DMSO-solubilized MDIVI-1 (Cayman Chemical Company, Ann Arbor, MI) by adding these solutions into the embryo water at 24 hpi. Larvae were exposed to DMSO, ramipril or MDIVI-1 for one hour at 33°C in a dark incubator. After exposure, zebrafish were transferred to 50 mL of fresh embryo water twice to rinse away the DMSO or DMSO-solubilized drugs. The final concentrations used for ramipril was 0.2 nM[21], and 7 nM for MDIVI-1[22].

### Survival Studies

For survival studies, mortality was monitored and counted daily in infected and control-injected larvae for up to 7 dpf. Larvae were maintained in dishes (∼50 larvae/dish) at 33°C with embryo water changes every two days.

### Viral Burden Assays

Tissue culture infectious dose 50 (TCID_50_) end-point dilution assays using MDCK/London cells were used to measure viral burden of influenza virus in zebrafish. Cohorts of 200 zebrafish per experimental group were used (see Zebrafish Care and Maintenance), infected (see Microinjections), and maintained in embryo water at 33°C in dishes (∼50 larvae/dish) with water changes every other day. Zebrafish were collected at 0, 24, 48, 72 and 96 hpi for larvae infected at 2 dpf, and 0, 24, 48 and 72 hpi for larvae infected at 3 dpf. At the appropriate timepoint, 25 larvae per group were collected and euthanized with an overdose (300 mg/L) of MS-222 for 10 minutes. Next, larvae were transferred into 500 µL of RNAlater (Invitrogen, Thermo Fisher Scientific), flash-frozen with liquid nitrogen, and stored at -80°C.

MDCK/London cells were plated into 96-well plates (PlateOne catalog number 1837-9600, USA Scientific) the evening before the TCID_50_ assay at a cell density (∼15,000 cells per well) to achieve 90-95% confluency the next day. After thawing the frozen samples on ice, the RNAlater was replaced with 500 µL of MEM-BSA-TPCK (see MDCK/London Cell Culture). Samples were homogenized using a Bullet Blender tissue homogenizer (Next Advance, Troy, NY) using a sterile metal bead at setting #3 for 5 minutes at 4°C and centrifuged at 8000 X g for 1 minute. Next, eight 1-to-8.5 serial dilutions (10^-0.9^ to 10^-7.4^) for each sample were prepared in MEM-BSA-TPCK. Cells were washed twice with PBS prior to adding the serial dilutions. After removing the second PBS wash, 50 µL serial dilutions for each sample were plated using triplicate wells (24 wells/sample), with 4 control wells per plate. 50 µL of MEM-BSA-TPCK was added to the control wells. Plates were centrifuged at 2000 × g for 10 minutes at 4°C and then incubated at 37°C for two hours with 5% CO_2_. Cell media was removed from all wells and 105 uL of MEM-BSA-TPCK was added to each well. Plates were then incubated at 37°C for 72 hours with 5% CO_2_. Next, cytopathic effects were observed, and cells counted using a Bio-Rad TC20 Automated Cell Counter (Bio-Rad, Hercules, CA). TCID_50_/ml was calculated using the Spearman-Kärber method[23].

### Respiratory Burst Assays

The capacity of zebrafish to generate ROS in vivo was quantified using a respiratory burst assay. Virus-infected and HBSS-injected zebrafish were collected at 24 and 48 hpi, anesthetized in sodium bicarbonate-buffered MS-222 (200 mg/L), and placed into black, flat bottom 96-well plates (Fluotrac 600, Greiner-Bio. Monroe, NC) with 100 µL of embryo water. HBSS-injected zebrafish were placed into the wells in the first 6 columns of the plate, and virus-infected zebrafish were placed into the remaining wells. The first 2 columns are treated with 5 µL of 1 mM protein kinase C inhibitor, bisindolylmaleimide I (BisI, Cayman Chemical Company), in DMSO. The BisI treated embryos were incubated for 30 minutes at 28°C. Zebrafish in columns 1, 3-4 and 7-9 were treated with 100 µL of embryo water plus 1 µg/mL 2′,7′-Dichlorofluorescein diacetate (H2DCFDA) (Sigma-Aldrich) in 0.4% DMSO. Zebrafish in columns 2, 5-6, and 10-12 were treated with 100 µL of embryo water plus 1 µg/mL H2DCFDA and 400 ng/ml phorbol 12-myristate 13-acetate (PMA, Sigma-Aldrich). The plates are covered with aluminum foil and incubated at 28°C for 2.5 hours. Fluorescence was read using a plate reader (BioTek Synergy, Agilent Technologies, Santa Clara, CA).

### Confocal Imaging

Zebrafish were anesthetized in sodium bicarbonate-buffered MS-222 (200 mg/L) and placed in a 24-well glass-bottom imaging plates (MatTek, Ashland, MA) with embryo water with 0.7% agarose. Olympus Fluoview IX-81 inverted microscope with FV1000 confocal system with 405, 458, 488, 514, and 543 nm laser lines was used for fluorescence and brightfield imaging of zebrafish. Images were obtained using x4 or x10 objectives.

### Image Analysis

Image analysis to count neutrophils and macrophages and quantify virus levels based on fluorescence intensities was conducted using MATLAB (version R2023a; The MathWorks Inc., Natick, MA). Longitudinal images obtained on the confocal were composed of 5 µm thin sections. Five sections proximal to the center of the zebrafish and five sections distal to the center of the zebrafish were analyzed for a total of 50 µm (approximately 40-50% of the total zebrafish width) was analyzed. Masks were generated to identify dsRed-tagged neutrophils, eGFP-tagged macrophages, and eCFP-PR8 Color-flu virus. Those masks were then used to count both neutrophils and macrophages, and quantify virus particles (pixels).

### Statistical Analysis

GraphPad Prism 9.0 (GraphPad Software, Boston, MA) was used to generate and analyze survival curves and graphs. Kaplan-Meier survival analysis was used to analyze survival curves with 95% confidence intervals. Mantel-Cox test p-values < 0.05 between sample groups were considered significant. Two-way ANOVA followed by Dunnett’s multiple comparison tests were used to analyze the Color-flu TCID_50_/mL values. A Brown-Forsythe one-way ANOVA test was used to analyze the TCID_50_/mL values from the live PR8, heat-inactivated and UV-inactivated larvae followed by Dunnett’s multiple comparison tests. Statistical analysis of the fold induction from respiratory burst assays were conducted using the Kruskal-Wallis test with the Dunn’s multiple comparison test for pairwise comparisons. Pairwise comparisons with an adjusted p-value < 0.05 were considered significant.

## RESULTS

### IAV Infection Decreases Survival and Replicates in Zebrafish

It was previously shown that zebrafish express α-2,6-linked sialic acid-containing receptors and PR8 H1N1 and X-31 A/Aichi/68 H3N2 IAV reduced survival of and replicated within zebrafish with disseminated infection[13]. We modified the original zebrafish IAV infection protocol to achieve approximately 50% mortality by 7 days post fertilization (dpf) by infecting embryos with PR8 IAV at either 2 or 3 dpi. In this protocol, we microinjected a 1 nL solution of 87% IAV and 13% diluent (sterile Hank’s Buffered Salt Solution (HBSS)) into the duct of Cuvier (DC) in 2 or 3 dpf anesthetized embryos. We found reduced survival in AB zebrafish systemically infected with three different PR8 virus lots at 2 dpf over vehicle (HBSS) controls (Figure 1A). The percent survival after 5 days was 57.6%, 55.6%, and 57.6% following infection with the three lots. Similar reductions in survival were observed in zebrafish systematically infected with two different PR8 lots at 3 dpf over vehicle controls (Figure 1B). After 4 days, 55.3% and 56.4% of larvae survived following injection by the two lots. Injection of heat- and ultraviolet (UV)-inactivated PR8 in 2 dpf AB zebrafish did not alter survival compared to vehicle controls (Supplemental Figure S1A). Increased viral titer was observed at 24 hpi in 2 dpf-infected AB zebrafish, but not in heat- and UV-inactivated PR8 (Supplemental Figure S1B).

**Figure 1:**
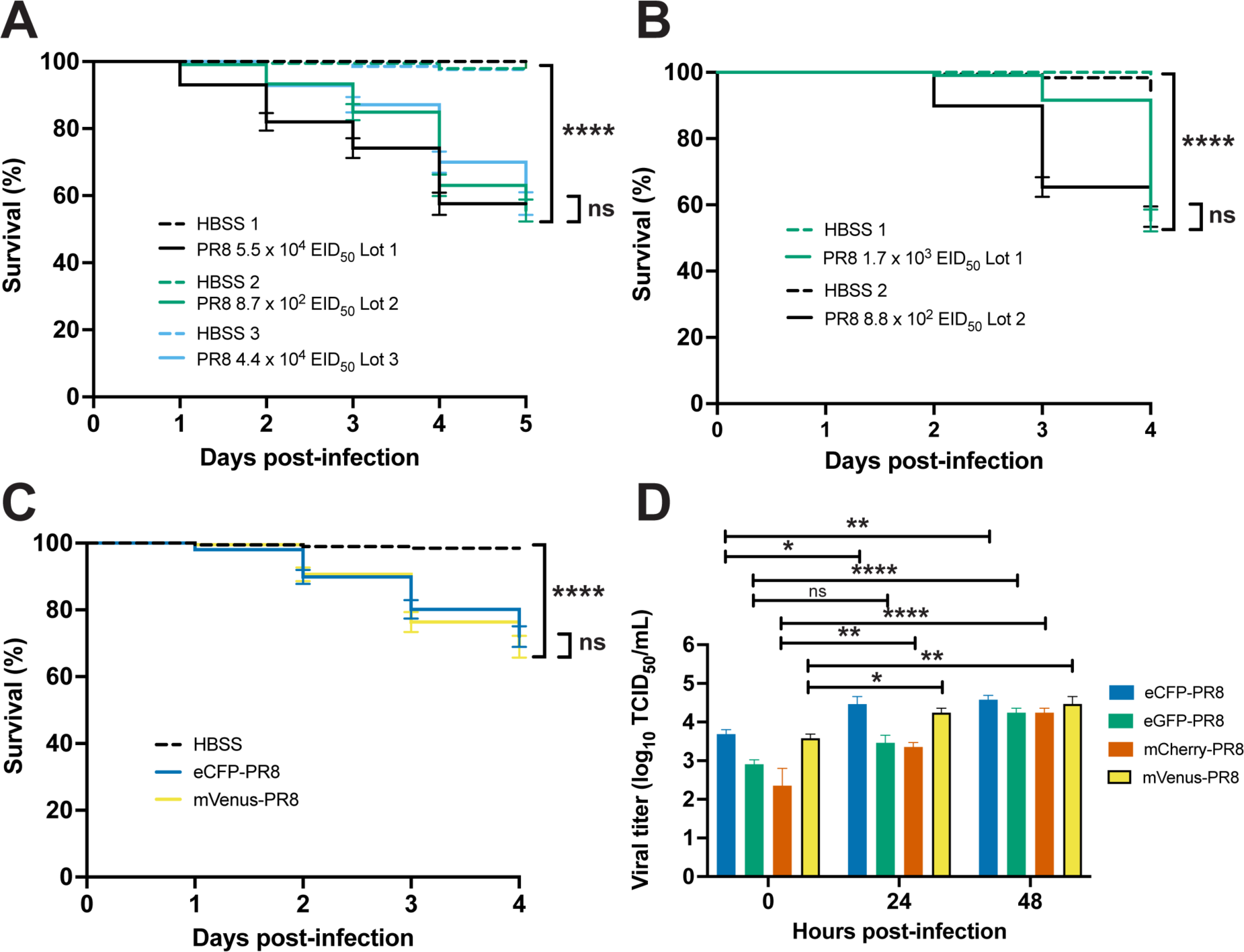
Characterization of PR8 and Color-flu systemically-infected AB zebrafish. A) Decreased survival of AB zebrafish systemically-infected with three different lots of PR8 IAV at 2 dpf compared to vehicle (HBSS) controls (p < 0.0001 for each lot comparison). Survival of PR8-infected embryos were not significantly different between lots (p = 0.0603). B) Decreased survival of AB zebrafish systemically-infected with two different lots of PR8 IAV at 3 dpf compared to vehicle controls (p < 0.0001 for each lot comparison). Survival of PR8-infected embryos were not significantly different between lots (p = 0.1442). C) Decreased survival of AB zebrafish systemically-infected with eCFP-PR8 or Venus-PR8 (3.2 × 10^2^ TCID50/mL) compared to vehicle controls (p < 0.0001 for each comparison). Survival of eCPF-PR8- or Venus-PR8-infected zebrafish were not significantly different (p = 0.5238). D) Increased TCID50 viral titer in Color-flu systemically-infected AB zebrafish at 24 and 48 hpi compared to 0 hpi. eCFP-PR8-, mCherry-PR8-, and Venus-PR8-infected zebrafish had increased viral titer at 24 and 48 hpi compared to 0 hpi (adjusted p-values = 0.0122 and 0.0044, respectively for eCFP-PR8; 0.0016 and < 0.001, respectively for mCherry-PR8; and 0.0321 and 0.0044, respectively for Venus-PR8). eGFP-PR8-infected zebrafish had increased viral titer at 48 hpi compared to 0 hpi (adjusted p-value < 0.0001).

### Color-flu Infection Decreases Survival and Replicates in Zebrafish

Using Color-flu[9] stocks kindly provided by Dr. Yoshihiro Kawaoka, we examined survival and viral burden in systemically infected zebrafish at 3 dpf (Figures 1C, 1D, S1C and S1D). Due to the lower virulence of Color-flu[9], our infection protocol was modified to inject a 6 nL solution of 87% Color-flu and 13% HBSS into the DC, and 6 nL of HBSS for controls. Decreased survival was observed in for all 4 strains (Figure S1D). Consistent with studies of the Color-flu strains in mice[9], we observed higher survival for zebrafish infected with Color-flu than PR8. Larvae survival after 4 days was lowest for the mVenus-PR8 (66.7%) and eCFP-PR8 (69.6%) strains and higher for the mCherry-PR8 (80.0%) and eGFP-PR8 (80.8%) strains. Consistent with the virulence shown in the survival studies, zebrafish systemically infected with Venus-PR8 and eCFP-PR8 had higher viral titers at 0 hpi than the other two Color-flu strains (Figure 1D). Significant increases in viral titers were observed at 24 hpi for all Color-flu strains except eGFP-PR8, and for all strains at 48 hpi. The expression of three proinflammatory genes was increased at 24 hpi with mVenus-PR8 infection at 24 hpi (Figure S2). Interferon regulatory factor 9 (*irf9*) activates the type I interferon response during viral infection[24]. Humans with mutations in *IRF9* have the immunologic disorder, Immunodeficiencey-65, and are susceptible to viral infections[24, 25]. Zebrafish receptor (TNFRSF)-interacting serine-threonine kinase 1, like (ripk1l) is orthologous to human RIPK1 which is participates in toll-like receptor (TLR) and retinoic acid-inducible gene I (RIG-I) signaling following viral infection[26]. Suppressor of cytokine signaling 3b (*socs3b*) is the zebrafish ortholog of human *SOCS3* and is a negative regulator of cytokine signaling and has been shown to be upregulated following influenza virus infection and result in overexpression of interleukin 6[27]. The expression of neutrophil cytosolic factor 1 (*ncf1*), which encodes a subunit of NADPH oxidase, was decreased at 24 hpi with mVenus-PR8 infection at 24 hpi (Figure S2).

### Zebrafish Lines Respond Differently to Influenza Infection

Multiple zebrafish lines have been used to model a wild-type response to injury and infection. To evaluate the consistency of the response to IAV infection, we compared the response of three lines to infection: AB, the wild line, EkkWill (EK), and the pigmentation mutant, *casper* (*mitfa^w2/w2^; mpv17^a9/a9^*). The AB line was originally used to establish the zebrafish as a model for studying the innate immune response to influenza virus[13] as well as bacterial[28] and fungal infection[29]. The EK line has been used to study fin[30] and cardiac[31] tissue regeneration. The *casper* line is optically transparent throughout development and into adulthood allowing for various studies including stem cell and tumor biology[14]. Systemic PR8 infection in 2 dpf embryos resulted in reduced survival for all three lines compared to HBSS vehicle controls (Figure 2A). *Casper* larvae had the lowest survival (45.7%) after 5 days, followed by EK (53.7%) and AB (62.7%). For embryos infected with PR8 at 3 dpf and followed for 4 days, *casper* larvae also had the lowest survival (44.2%) compared with EK (63.0%) and AB (62.6%) (Figure 2B).

**Figure 2:**
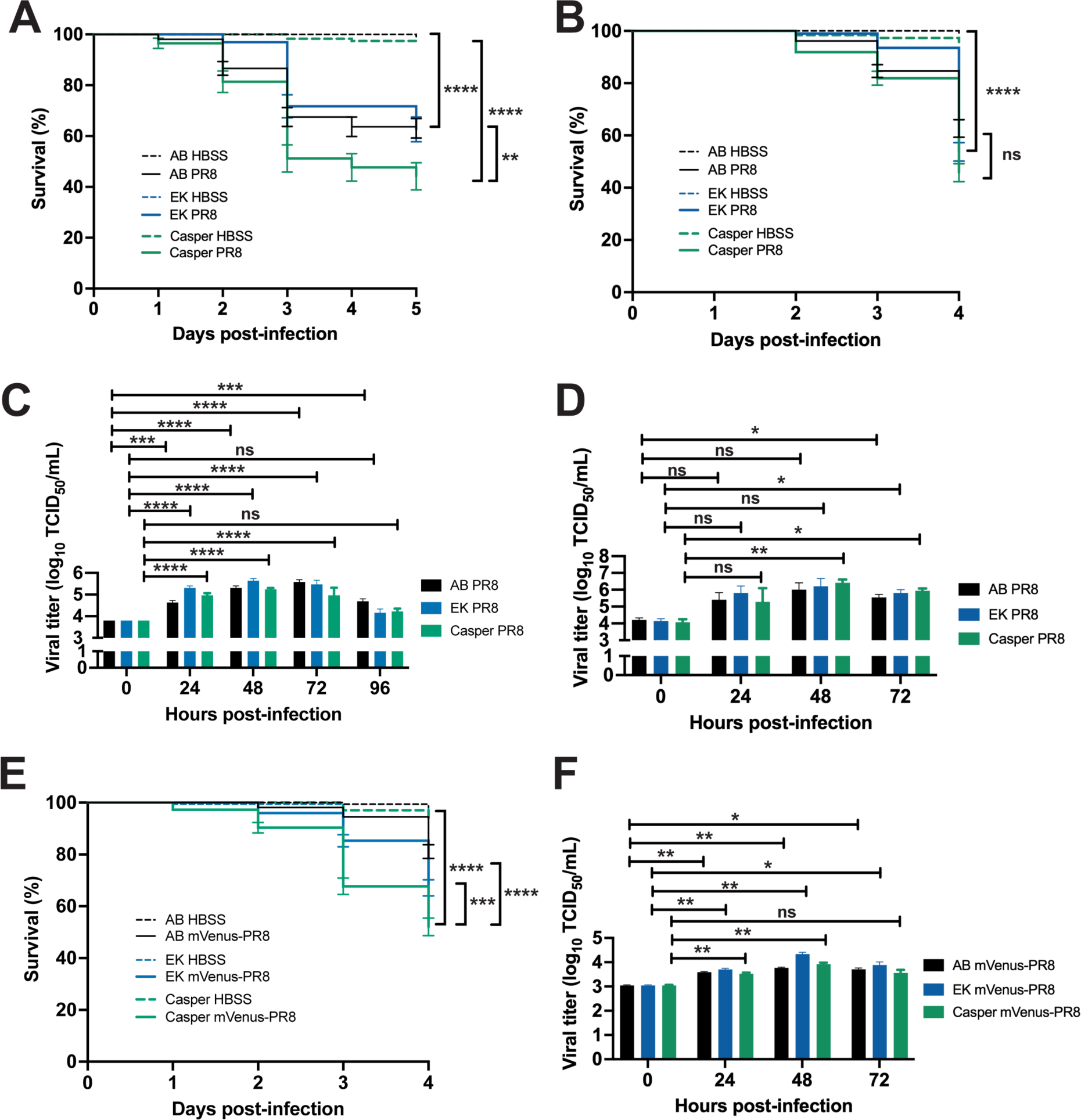
Characterization of PR8 and Color-flu systemically infected across zebrafish lines. A) Decreased survival was greater in *casper* than AB or EK lines with 2 dpf systemic PR8 infection. PR8 infected zebrafish had decreased survival over vehicle (HBSS) controls (p < 0.0001 for all lines). *Casper* zebrafish had lower survival than EK (p = 0.0024) and AB (p = 0.0055) by log-rank Mantel-Cox test. B) Decreased survival was observed in AB, EK and *casper* with 3 dpf -PR8 infection over - controls (p < 0.0001 for all lines). No significant (ns) difference was detected in survival between PR8-infected AB, EK and *casper* larvae. C) Increased TCID50 viral titer in 2 dpf PR8 infected AB, EK and *casper* zebrafish at 24, 48 and 72 hpi compared to 0 hpi (adj. p-value < 0.001 for all comparisons except for AB 24 hpi (adj. p-value = 0.0006), and AB 96 hpi (adj. p-value = 0.0003)). D) Increased TCID50 viral titer in 3 dpf PR8 infected AB, EK and *casper* zebrafish at 72 hpi compared to 0 hpi (adj. p-value = 0.0323, 0.0211, and 0.0452, respectively), and *casper* zebrafish at 48 hpi compared to 0 hpi (adj. p-value = 0.0063). E) Decreased survival with 3 dpf mVenus-PR8 infected AB, EK and *casper* zebrafish compared to HBSS controls (p < 0.0001 for all lines). As observed with PR8 infection, *casper* zebrafish had the lowest survival (67.7%) that was lower than AB (94.5%, p < 0.0001) and EK (85.5%, p < 0.0002). F) eCFP-PR8-, mCherry-PR8-, and Venus-PR8-infected zebrafish had increased viral titer at 24 and 48 hpi compared to 0 hpi (adjusted p-values = 0.0122 and 0.0044, respectively for eCFP-PR8; 0.0016 and < 0.001, respectively for mCherry-PR8; and 0.0321 and 0.0044, respectively for Venus-PR8). eGFP-PR8-infected zebrafish had increased viral titer at 48 hpi compared to 0 hpi (adjusted p-value < 0.0001).

Viral load was measured at daily intervals following systemic PR8 infection of 2 dpf embryos from all three lines using tissue culture infectious dose (TCID_50_) virus titer assays (Figure 2C). Viral titers increased by 24 hpi in all three lines, peaked at 48-72 hpi, and then declined by 96 hpi. AB larvae had peak viral load at 72 hpi and still had increased load at 96 hpi. EK and *casper* larvae had peak viral load at 48 hpi with levels not different from the 0 hpi controls by 96 hpi. Viral load also increased with system PR8 infection in 3 dpf embryos from the three lines by 72 hpi (Figure 2D). *Casper* larvae had peak viral load at 48 hpi like the 2 dpf infected embryos.

Localized PR8 infection in the swimbladder of 4 dpf AB, EK and *casper* larvae also resulted in decreased survival by 3 dpi over HBSS controls (Supplemental Figure S2A). Like the survival observed with systemic infection, PR8-infected *casper* larvae also had the lowest survival (31.5%) compared to EK (54.5%) and AB (61.9%). Likewise, localized Color-flu (mVenus-PR8) infection in the swimbladder of 4 dpf AB, EK and *casper* larvae also resulted in reduced survival with *casper* having the lowest percent survival (42.3%) followed by EK (67.5%) and AB (74.1%) (Supplemental Figure S2B).

### Live Confocal Imaging of Zebrafish Infected with Color-flu

Optical transparency of zebrafish embryos and larvae allow for *in vivo* confocal imaging of Color-flu where host-pathogen interactions can be visualized. With disseminated infection, virus was observed throughout AB larvae for each of the four Color-flu strains (Figure 3). In these whole larvae lateral views at 24 hpi, the virus accumulated highest in the yolk sac and yolk sac extension. The virus was also observed in the head and skeletal muscle. Several zebrafish transgenic fluorescent reporter strains have been used to visualize cell types, including neutrophils and macrophages. We injected the dual macrophage and neutrophil reporter line, Tg(*mpeg1:*eGFP*;lyz:*dsRed), with eCFP-PR8 and vehicle (HBSS) control at 3 dpf and imaged larvae at 24 hpi (Figure 3B). Control larvae show macrophages and neutrophils in the circulatory system and other tissues including skeletal muscle. Imaging of eCFP-PR8 infected larvae allow for simultaneous visualization of macrophages, neutrophils, and virus particles.

**Figure 3:**
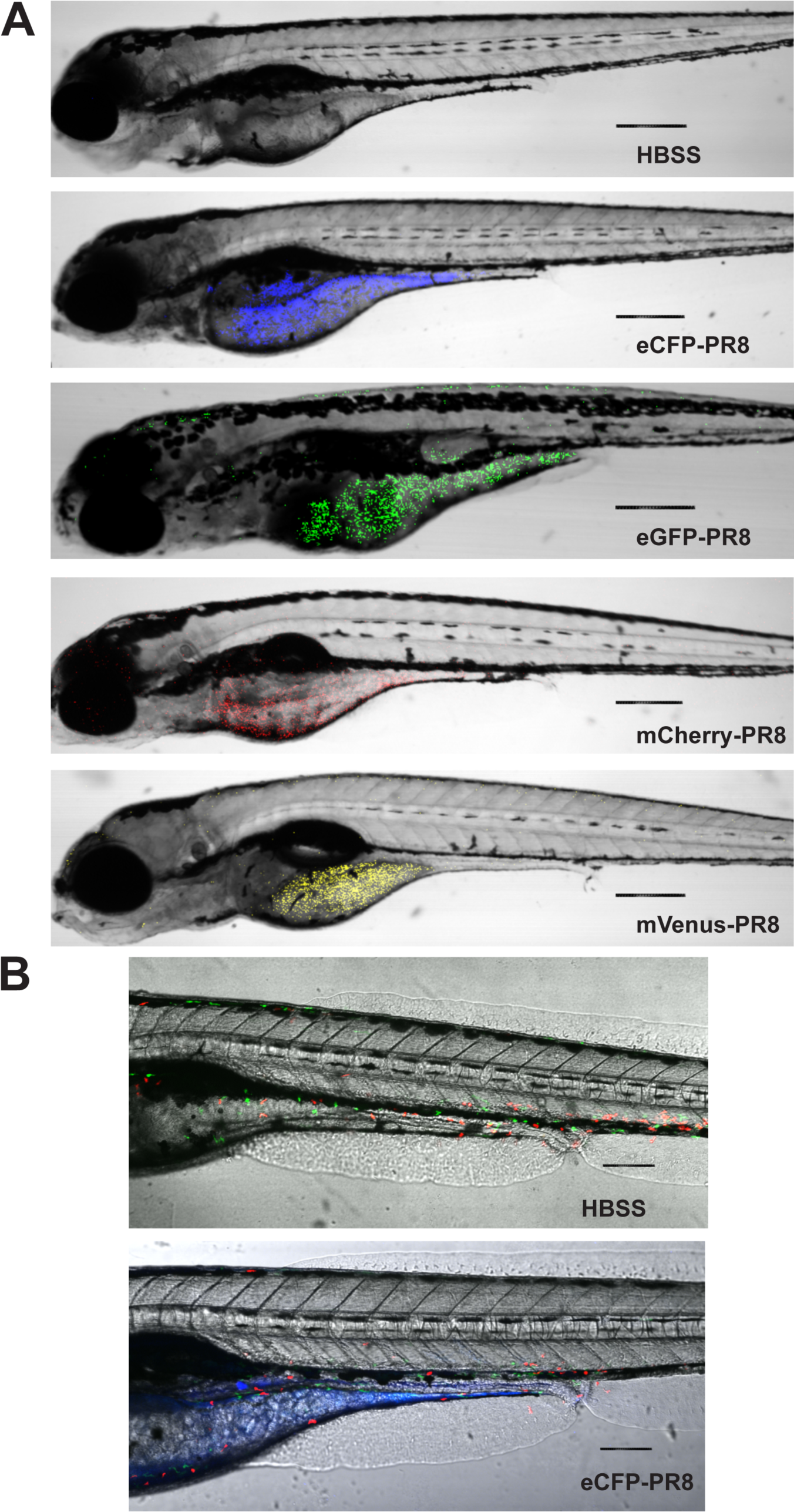
Confocal imaging of Color-flu infected zebrafish. A) Larvae at 24 hours post systemic injection of HBSS, eCFP-PR8, eGFP-PR8, mCherry-PR8, and mVenus-PR8 at 3 dpf at 4× resolution. Scale bar: 300 mm. B) Tg*(mpeg1:e*GF*P;lyz:*dsRed) larvae at 24 hours post systemic injection of HBSS, eCFP-PR8 at 3 dpf at 10x resolution showing neutrophils (red), macrophages (green) and eCFP-PR8 (blue). Scale bar: 700 um.

### Evaluating Small Molecules That Alter the Response to IAV Infection

Next, we evaluated how two small molecule drugs, ramipril and MDIVI-1, would alter the response to IAV infection by examining survival, virus burden, immune capacity by respiratory burst assay, and relative immune cell proliferation and virus load using confocal imaging. To model drug therapies administered after infection, embryos were infected with IAV at 3 dpf, and then treated with either DMSO (control), ramipril, or MDIVI-1 at 24 hpi for one hour, and then washed and maintained in clean embryo water for up to 4 days post infection. Ramipril or MDIVI-1 administered to PR8-infected and vehicle (HBSS) control larvae were at a concentration of 0.2 nM or 7nM, respectively, increased survival (Figure 4). Ramipril increased survival to 90.0% compared to 52.1% in DMSO controls. MDIVI-1 increased survival to 85.4% compared to 55.4% in DMSO controls. With mVenus-PR8 infected larvae, ramipril or MDIVI-1 treatment increased survival to 87.7% and 85.0%, respectively, compared to 58.9% in DMSO controls. The ramipril and MDIVI-1 concentrations used were selected after evaluating differences in survival due to drug treatment were evaluated for a range of doses for ramipril (0.1, 0.2, 0.3 and 0.4 nM) and MDIVI-1 (3, 5, 7 and 10 nM) in PR8-infected larvae at either 2 or 3 dpf (Figures S3 and S4). Both 0.1 and 0.2 ramipril treatment, and only 7 nM MDIVI-1 increased survival in larvae infected with PR8 at both 2 and 3 dpf. The highest doses of ramipril (0.4 nM) and MDIVI-1 (10 nM) both decreased survival in control (HBSS) injected larvae.

**Figure 4:**
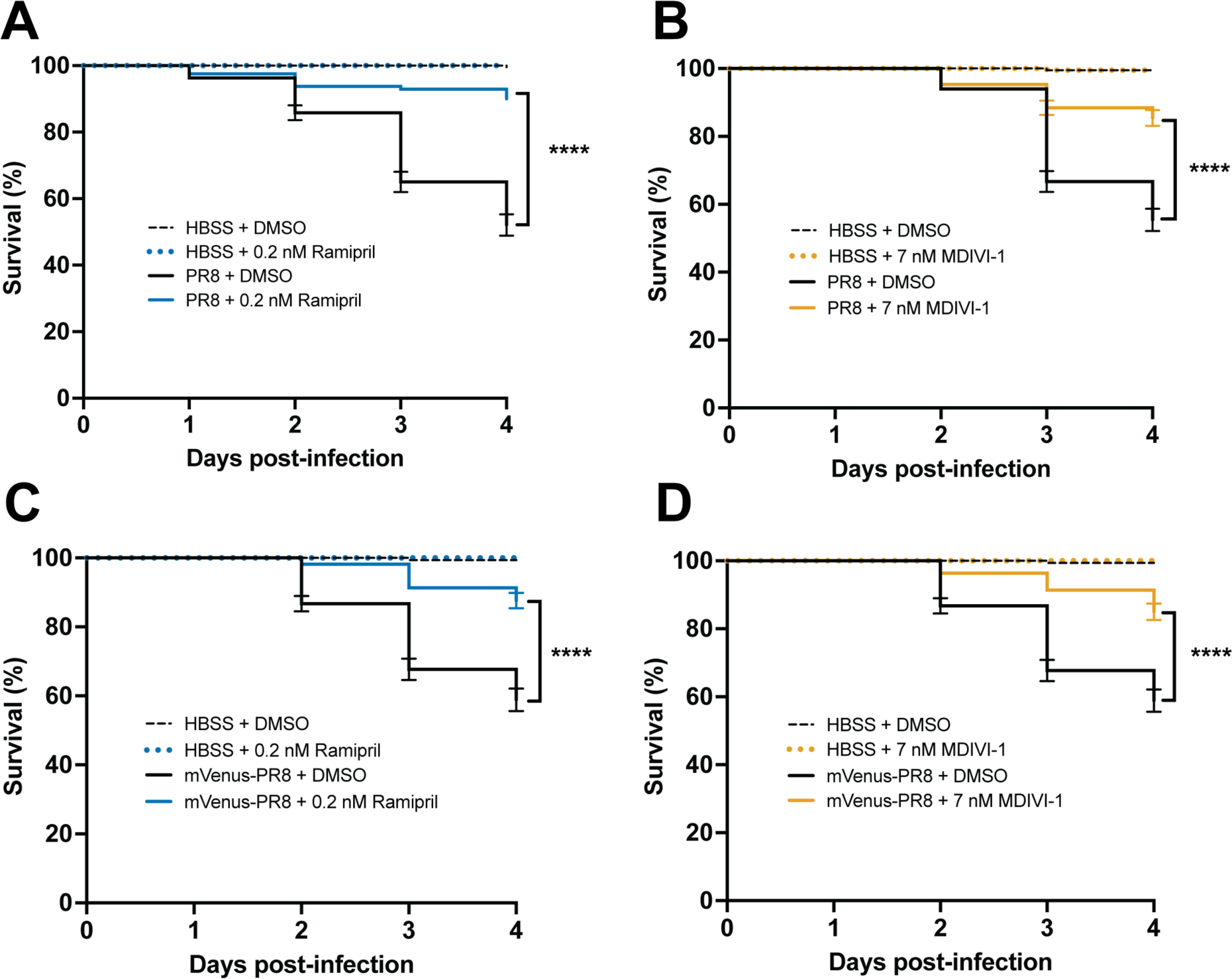
Ramipril and MDIVI-1 increase survival in systemically infected PR8 and Color-flu infected 3 dpf zebrafish. A) Increased survival in PR8 infected AB zebrafish treated with 0.2 nM ramipril compared to DMSO controls (p < 0.0001). B) Increased survival in PR8 infected AB zebrafish treated with 7 nM MDIVI-1 compared to DMSO controls (p < 0.0001). C) Increased survival in mVenus-PR8 infected AB zebrafish treated with 0.2 nM ramipril compared to DMSO controls (p < 0.0001). D) Increased survival in mVenus-PR8 infected AB zebrafish treated with 7 nM MDIVI-1 compared to DMSO controls (p < 0.0001). As the survival studies shown in A-D were conducted simultaneously, the data are represented in separate panels to make the data legible. As such, the data for some groups are shown in multiple panels as follows: 1) the same DMSO HBSS and DMSO PR8 data are shown in A-D; 2) the same 0.2 nM rampiril HBSS data are shown in A and C; and 3) the same 7 nM MDIVI-1 HBSS data are shown in B and D.

Ramipril and MDIVI-1 treatment reduced viral burden in IAV-infected larvae by 24 hours after treatment (Figure 5). In larvae infected with PR8 at 2 and 3 dpf, viral burden increased at 24 hpi when larvae were treated with ramipril, MDIVI-1 or DMSO. By 48 hpi, viral burden was reduced in ramipril and MDIVI-1 treated larvae. In the 2 dpf infected larvae with ramipril and MDIVI-1 treatment, viral burden at 72 hpi was reduced to levels measured at 0 hpi. A reduction in viral burden following ramipril and MDIVI-1 treatment was also observed at 48 hpi with mVenus-PR8 infection (Figure 5C).

**Figure 5:**
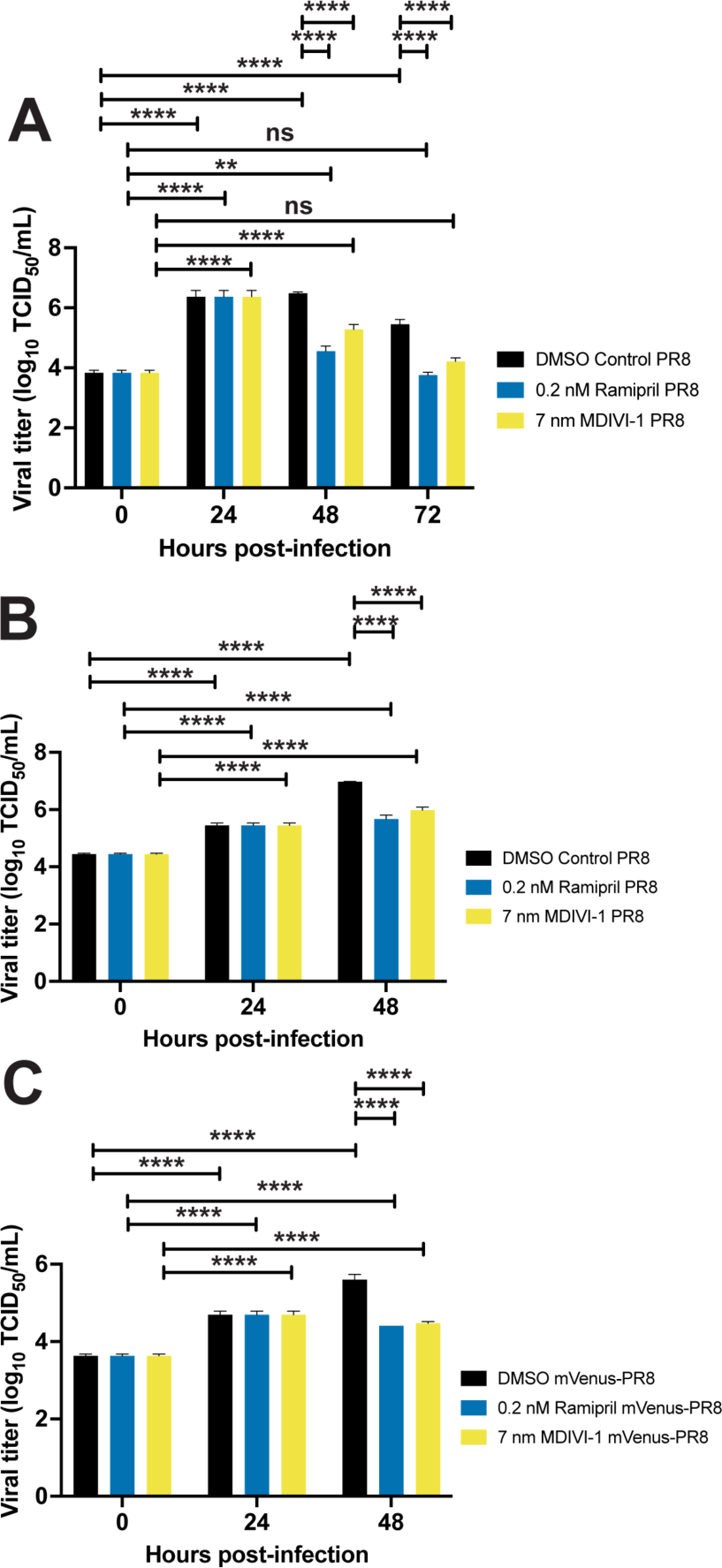
Ramipril and MDIVI-1 treatment lower viral burden in IAV infected zebrafish. A) Viral titers for both ramipril (0.2 nM) and MDIVI-1 (7 nM) treated 2 dpf zebrafish infected with PR8 were significantly higher from 0 hpi at 24 and 48 hpi, but not different at 72 hpi. For DMSO-treated zebrafish, viral titers were significantly increased at 24, 48 and 72 hpi (p < 0.0001 for each comparison). For ramipril-treated zebrafish, viral titers were significantly increased at 24 (p < 0.0001) and 48 hpi (p = 0.0022). For MDIVI-1-treated zebrafish, viral titers were also significantly increased at 24 and 48 hpi (p < 0.0001 for each comparison). B) Viral titers for ramipiril and MDIVI-1 treated 3 dpf zebrafish infected with PR8 were significantly lower by 48 hpi (p < 0.0001 for each comparison). C) Viral titers for ramipril and MDIVI-1 treated 3 dpf zebrafish infected with mVenus-PR8 were significantly lower at 48 hpi (p < 00001 for each comparison).

Next, the level of ROS generated by larvae following infection was quantified to characterize the capacity of the immune system to mount a respiratory burst response. In PR8- and mVenus-PR8-infected larvae, the respiratory burst response was reduced compared to uninfected controls (Figure 6). Ramipril and MDIVI-1 treatment rescued the response such that the level of induction was the same or higher than the uninfected controls.

**Figure 6:**
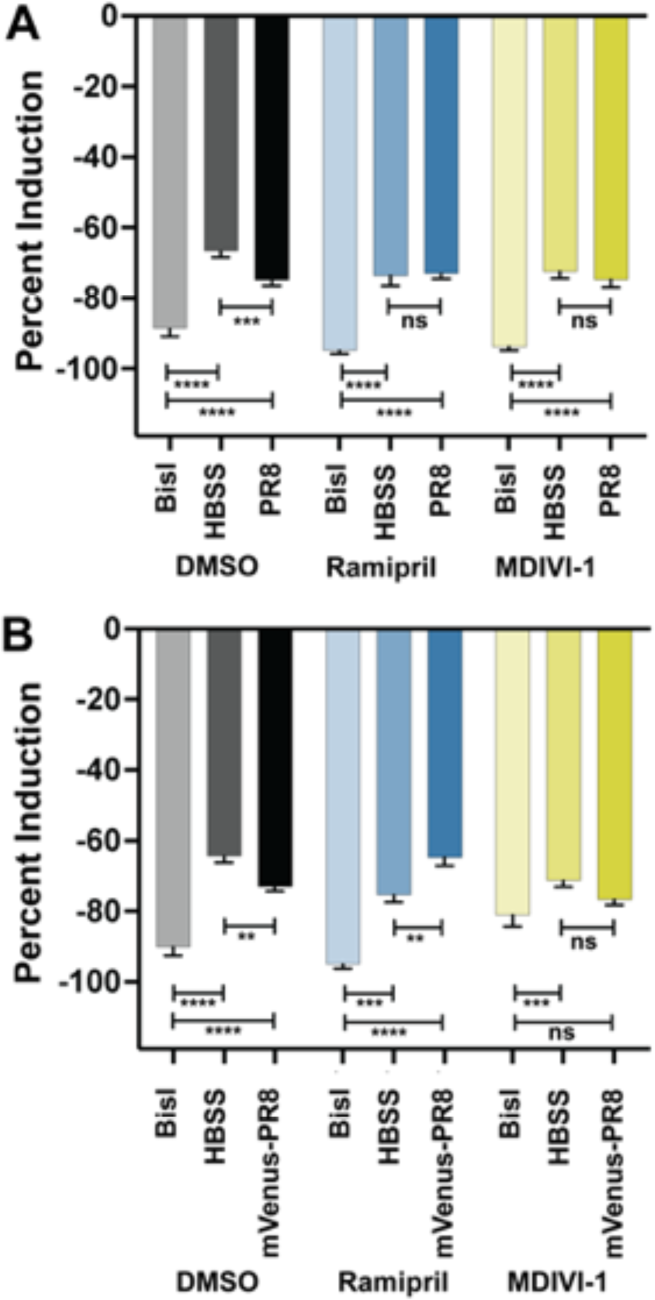
Ramipril and MDIVI-1 treatment alters respiratory burst response in IAV systemically infected zebrafish at 48 hpi. A) Respiratory burst response in larvae treated with DMSO, ramipril (0.2 nM), and MDIVI-1 (7 nM) at 48 hpi following systemic injection at 3 dpf with PR8 or HBSS. PR8 infection decreased the response over HBSS in DMSO-treated controls (adjusted p = 0.0008). Both ramipril and MDIVI-1 treatment rescued the reduction in respiratory burst response as PR8-infected larvae had the same response as HBSS injected controls (comparisons were not significant (ns)). The protein kinase C inhibitor, bisindolylmaleimide I (BisI), was used as a positive control as it suppresses the respiratory burst response (adj. p < 0.0001 for all comparisons). B) Respiratory burst response in larvae treated with DMSO, ramipril (0.2 nM), and MDIVI-1 (7 nM) at 48 hpi following systemic injection at 3 dpf with mVenus-PR8 or HBSS. Similar to the PR8-infected larvae, mVenus-PR8 infection decreased the response over HBSS in DMSO-treated controls (adj. p = 0.0040). Ramipril treatment resulted in a higher respiratory burst response with mVenus-PR8 infection than HBSS controls (adj. p = 0.0049). MDIVI-1 treatment rescued the reduction in respiratory burst response as mVenus-PR8 infected larvae had the same response as HBSS injected controls. The BisI controls were different for the DMSO (adj. p < 0.0001 for both), ramipril (adj. p = 0.0003 for HBSS, and adj. p < 0.0001 for mVenus-PR8), and MDIVI-1 (adj. p = 0.0006 for HBSS, ns for mVenus-PR8) groups.

We then characterized the abundance of neutrophils, macrophages and virus particles using fluorescent confocal imaging of Tg(*mpeg1:eGFP;lyz:dsRed*) larvae infected with eCFP-PR8. In DMSO treated larvae we observed higher numbers of neutrophils and virus particles, but lower numbers of macrophages at 48 hpi (Figure 7). Ramipril increased the number of neutrophils in infected larvae compared to uninfected controls, a difference not observed with macrophages. With eCFP-PR8 infection, the number of macrophages were increased with MDIVI-1 treatment over DMSO controls. The number of virus particles in infected larvae were reduced with ramipril and MDIVI-1 treatment compared to DMSO controls.

**Figure 7:**
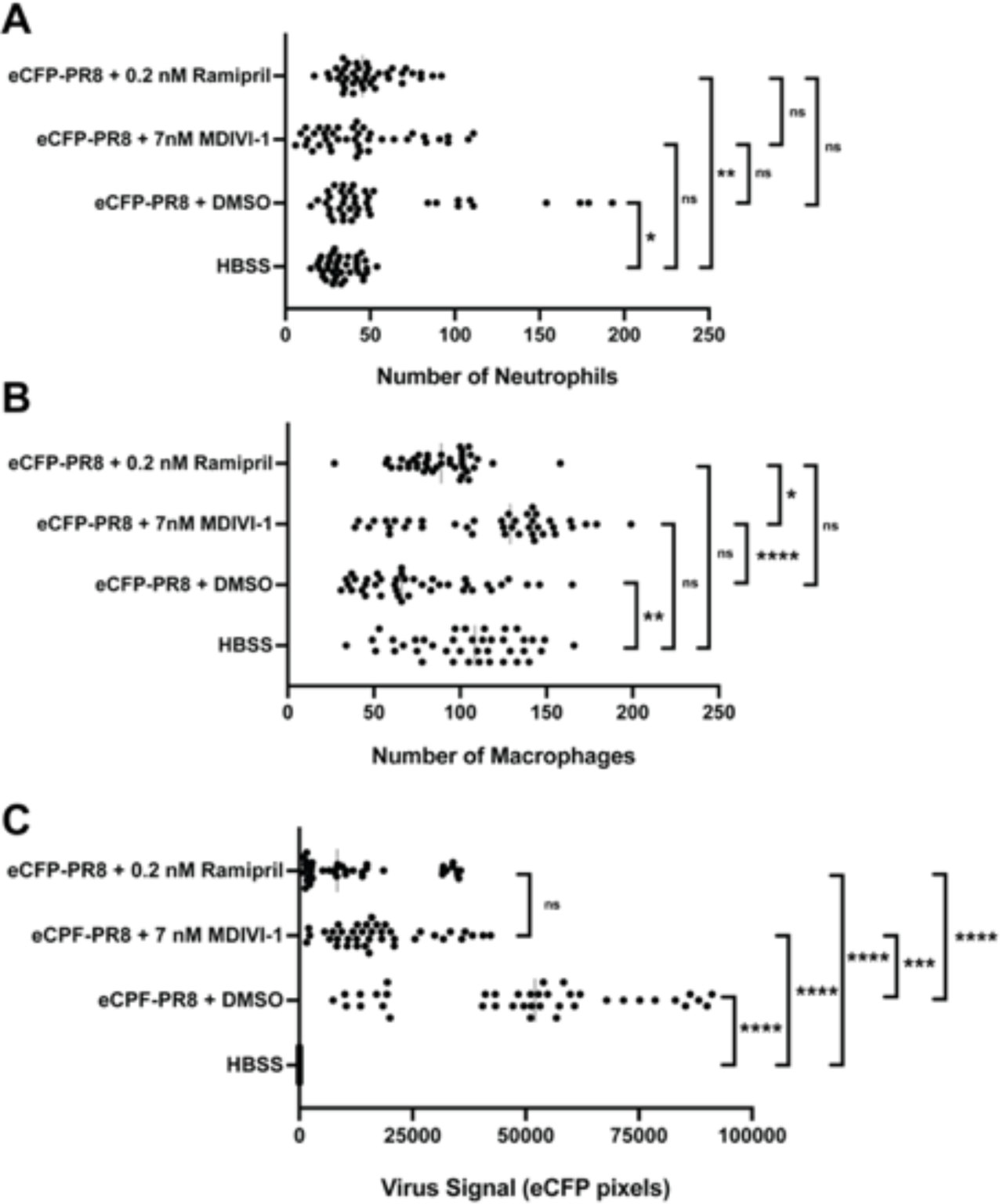
Quantification of neutrophils (A), macrophages (B), and level of virus (C) by fluorescent confocal imaging of Tg*(mpeg1:*eGF*P;lyz:*dsRed) larvae at 48 hours post injection by eCFP-PR8 or HBSS following treatment by DMSO, ramipril and MDIVI-1. A) Number of neutrophils were increased with eCFP-PR8 infection following DMSO treatment over HBSS controls (adjusted p-value = 0.0145), and with ramipril treatment (adj. p-value = 0.0010), but not with MDIVI-1 treatment. B) The number of macrophages were decreased in eCFP-PR8 infected larvae treated with DMSO over HBSS controls (adj. p-value = 0.0020), but was not different with ramipril or MDIVI-1 treatment. MDIVI-1 treatment increased the number of macrophages over DMSO controls (adj. p-value < 0.0001), and over ramipril treated larvae (adj. p-value = 0.0126). C) The level of virus was higher in eCFP-PR8 infected larvae treated with DMSO, ramipril, and MDIVI-1 (adj. p-value < 0.0001 for all comparisons). The level of virus was lower with ramipril (adj. p-value < 0.0001) and MDIVI-1 (adj. p-value = 0.0008) treatment over the DMSO-treated controls.

Time-lapse *in viv*o confocal imaging allows cells to be tracked in these transgenic lines injected with Color-flu and then exposed to small molecules. Time-lapse imaging showed circulating neutrophils and macrophages over the course of 30 minutes in a 5 dpf larvae that were injected with HBSS at 3dpf (Supplemental Movie 1). In a 5 dpf larvae infected with eCFP-PR8 at 3 dpf and then exposed to DMSO at 24 hpi for one hour, we observed virus particles in several tissues, both macrophages and neutrophils near the highest density of virus particles, and more circulating macrophages over the course of 30 minutes (Supplemental Movie 2). In eCFP-PR8-infected larvae exposed to ramipril, we observe less virus particles and more neutrophils and fewer macrophages over 30 minutes (Supplemental Movie 3). The number of macrophages were increased in eCFP-PR8-infected larvae exposed to MDIVI-1 over 30 minutes (Supplemental Movie 4).

## DISCUSSION

Here, we demonstrate that Color-flu can be used to model the innate immune response to IAV infection in zebrafish larvae, and that both ramipril and MDIVI-1 treatment improve survival and reduce viral burden following IAV infection. There is an urgent need to develop new antiviral therapies for IAV due to the threat of new IAV strains pose in the future. The IAV strains that caused the last four influenza pandemics since 1900 were derived from viral transmission of strains from avian and non-human mammalian hosts and subsequent recombination with human strains[32]. Furthermore, antiviral therapies can become ineffective when the right combination of mutations occur in the virus. For example, a single amino acid substitution (H274Y) in neuraminidase was reported in H5N1 isolates that conferred resistance to oseltamivir[33]. The zebrafish model of IAV infection is complementary to other animal and cell line models used to evaluate antiviral therapies. We show how the model can be used to evaluate differences in survival, viral burden, respiratory burst, gene expression, and fluorescent confocal imaging. The zebrafish Color-flu model of IAV infection is unique as it is the only model where interactions between virus particles and host cells in an intact vertebrate can be visualized *in vivo* using fluorescent confocal imaging. As zebrafish larvae only have innate immune cells at this stage of development, the model can be used to study the roles of neutrophils and macrophages during IAV infection. Studies of the innate immune response to IAV is needed as the virus can evade the innate response through interactions of viral proteins (PB1, PB1-F2, PB2, PA, and NS1) with the host[34–39].

We demonstrate that zebrafish larvae are a robust model for the innate immune response to IAV infection by evaluating multiple lots of PR8 virus, and multiple zebrafish lines. The landmark paper that first established the zebrafish larvae as a model for IAV infection[13] had studied the AB line. Here, we demonstrate infection in two “wild-type” lines, AB and EK, along with the pigmentation mutant, *casper*. When the same amount of virus is injected in larvae from these strains, *casper* had the lowest level of survival and highest virus burden. As done in other model organisms, comparing strains with varying phenotypes can be a powerful way to understand disease mechanisms[40]. Whole genome sequencing of *casper*, a related pigmentation mutant, *roy*, and AB zebrafish revealed 4.3 million single nucleotide polymorphisms between strains, including the mutations in the *mpv7* and *mitfa* genes in the *casper* line[41]. The combination of sequence variants found in *casper* could potentially explain the difference in response to IAV infection.

Visualizing the interaction between fluorescently labeled virus particles and host immune cells in living larvae makes the zebrafish Color-flu model unique. Zebrafish larvae from green or red fluorescent reporter strains that label neutrophils and/or macrophages have enabled studies of host-pathogen interactions, such as those during bacterial and fungal infections. However, many of these studies have often included the use of a green or red fluorescently labeled pathogen thereby preventing the simultaneous imaging of the pathogen with both host immune cell types. As we demonstrate, the four Color-flu strains developed by the Kawaoka laboratory makes it possible to image virus particles along with neutrophils and macrophages using distinct color combinations. Furthermore, the Color-flu strains have retained their virulence despite the introduction of the fluorescent transgene as we demonstrate by showing decreased survival and increased viral burden in infected AB, EK and *casper* larvae.

We describe the response to both systemic and localized IAV infection. To study a systemic response, IAV was directly injected into the duct of Cuvier so that the virus can spread throughout the larvae via the circulatory system. Confocal imaging of Color-flu infected larvae show virus throughout different tissues of the larvae. Virus particles from Color-flu strains accumulate in the yolk sac and yolk sac extension. The yolk sac accumulation of compounds has been noted in toxicological studies using zebrafish larvae[42]. In a study of nanoplastics toxicity, embryos were exposed to 70 nm diameter fluorescent polystyrene nanoparticles in embryo water[43]. By 1 dpf, fluorescence was detected in the yolk sac and yolk sac extension of 1 and 10 ppm exposed embryos that persisted until 5 dpf when the yolk sac is reabsorbed after the gastrointestinal system becomes active. As the diameter of the nanoparticles were slightly smaller than IAV particles (80-120 nm), it is plausible that the virus accumulates in the yolk sac and yolk sac extension like these nanoparticles.

We applied the zebrafish Color-flu model to evaluate whether the ACE inhibitor, ramipril, or the autophagy inhibitor, MDIVI-1, would alter the response to IAV infection. Both small molecules improved the response to infection as treated larvae had increased survival, lower viral burden, and a respiratory burst response that was the same or higher than the uninfected controls. Quantification of fluorescent confocal images of Tg(*mpeg1:eGFP;lyz:dsRed*) larvae infected with Color-flu showed that the relative number of virus particles were lower in the ramipril and MDVI-1 treated larvae. Ramipril treatment increased the relative number of neutrophils in infected larvae whereas MDIVI-1 increased the relative number of macrophages.

Ramipril is an ACE inhibitor that is frequently prescribed for hypertension as ACE produces angiotensin II which, in turn, elevates blood pressure[44]. ACE has also been associated with innate immune function[45]. Angiotensin II mediates proinflammatory responses, including the production of reactive oxygen species (ROS) by NADPH oxidase and subsequent activation of nuclear factor-κB (NF-κB) and activator protein 1 (AP1)[46]. Overexpression of ACE expression in mouse neutrophils increased superoxide production and enhanced clearance of bacteria in mice infected with methicillin-resistant *Staphylococcus aureus* (MRSA), *Pseudomonas aeruginosa*, or *Klebsiella pneumoniae*[47]. ACE overexpression in mouse macrophages also lowered bacterial burden in mice infected with MRSA or *Listeria monocytogenes* [PMID: 20937811]. While ACE overexpression in myeloid cells benefit immune responses, ACE deficiency can also be beneficial. A study of mouse models of acute respiratory distress syndrome showed that ACE deficient mice have less lung damage[48]. Interestingly, a study of electronic health records of patients in the Clinical Practice Research Datalink in the United Kingdom showed a lower risk of influenza infection depending on the duration of ACE inhibitor use[17]. Our studies of ramipril treatment in our zebrafish IAV model demonstrated an improved response to viral infection.

Mitochondria have important roles in immune function, including regulating inflammation[49]. Mitophagy maintains mitochondrial function by clearing mitochondria when they become damaged[50]. A myriad of biological processes is altered when mitochondrial function is disrupted, including the overproduction of mitochondrial ROS that can activate the nucleotide oligomerization domain (NOD)-like receptor family pyrin domain containing 3 (NLRP3) inflammasome[51]. IAV infection has been shown to induce mitophagy thereby altering inflammasome activation[52, 53]. IAV nucleoprotein (NP) was shown to induce mitophagy through toll interacting protein (TOLLIP) and mitochondria antiviral signaling protein (MAVS)[54]. In that study, the mitophagy inhibitor MDIVI-1 was demonstrated to reduce NP-induced degradation of mitochondrial proteins. The mitochondrial fission inhibitor, MDIVI-1, was also shown to reduce NLRP3 inflammasome activation in transmitochondrial cybrid cells with diminished mitochondrial function[55]. In our studies of IAV-infected zebrafish larvae, we show that inhibiting mitophagy using MDIVI-1 treatment increases survival and decreases viral burden. We can therefore hypothesize that MDVI-1 counters IAV-induced mitophagy and NLRP3 inflammasome activation.

Zebrafish embryos and larvae have been used screen small molecule drugs where the response of separate larvae to different treatments can be monitored in multi-well plates[56]. Large-scale screening of small molecules that alter the innate immune response to IAV infection is feasible using our Color-flu model. Color-flu infected larvae would be exposed to small molecules within wells, and then characterized using fluorescent confocal imaging to determine relative abundance of virus particles. In these studies, small molecules could be evaluated using different concentrations and in also different combinations.

In summary, we demonstrate that Color-flu can be used to study the innate immune response to IAV infection in zebrafish larvae. This model is complementary to other models of IAV infection. The model is the only IAV model that allows for the visualization of interactions between virus particles and host cells in an intact vertebrate *in vivo*. The roles of neutrophils and macrophages during IAV infection can also be readily characterized using this model. Using Color-flu, we characterize how ACE inhibition by ramipril and mitophagy inhibition by MDIVI-1 both improve the response to IAV infection by limiting inflammation through distinct mechanisms. These studies demonstrate how larger small molecule screens are possible using this zebrafish Color-flu model of IAV infection. Such screening studies are needed to find small molecules that could be developed into novel antiviral therapies for influenza virus.

## ACKNOWLEDGEMENTS

We thank Dr. Yoshihiro Kawaoka of the University of Wisconsin for sharing the Color-Flu IAV strains, and Mark Nilan at the University of Maine Zebrafish Facility for excellent zebrafish care. This work was supported by the National Institute of Allergy and Infectious Diseases of the National Institutes of Health under grant number R15 AI131202, and Institutional Development Award (IDeA) from the National Institute of General Medical Sciences of the National Institutes of Health under grant numbers P20 GM144265 and P20 GM103423. These funding sources played no role in the design of the study, in collection, analysis, and interpretation of data, or in writing the manuscript.

## AUTHOR CONTRIBUTIONS

B.S. and B.K. designed the experiments and wrote the paper. B.S., A.B., H.F., J.G. and C.A. performed experiments. B.S., A.B., H.F., J.G. and B.K. analyzed the data.

## SUPPLEMENTAL FIGURES

**Figure S1:**
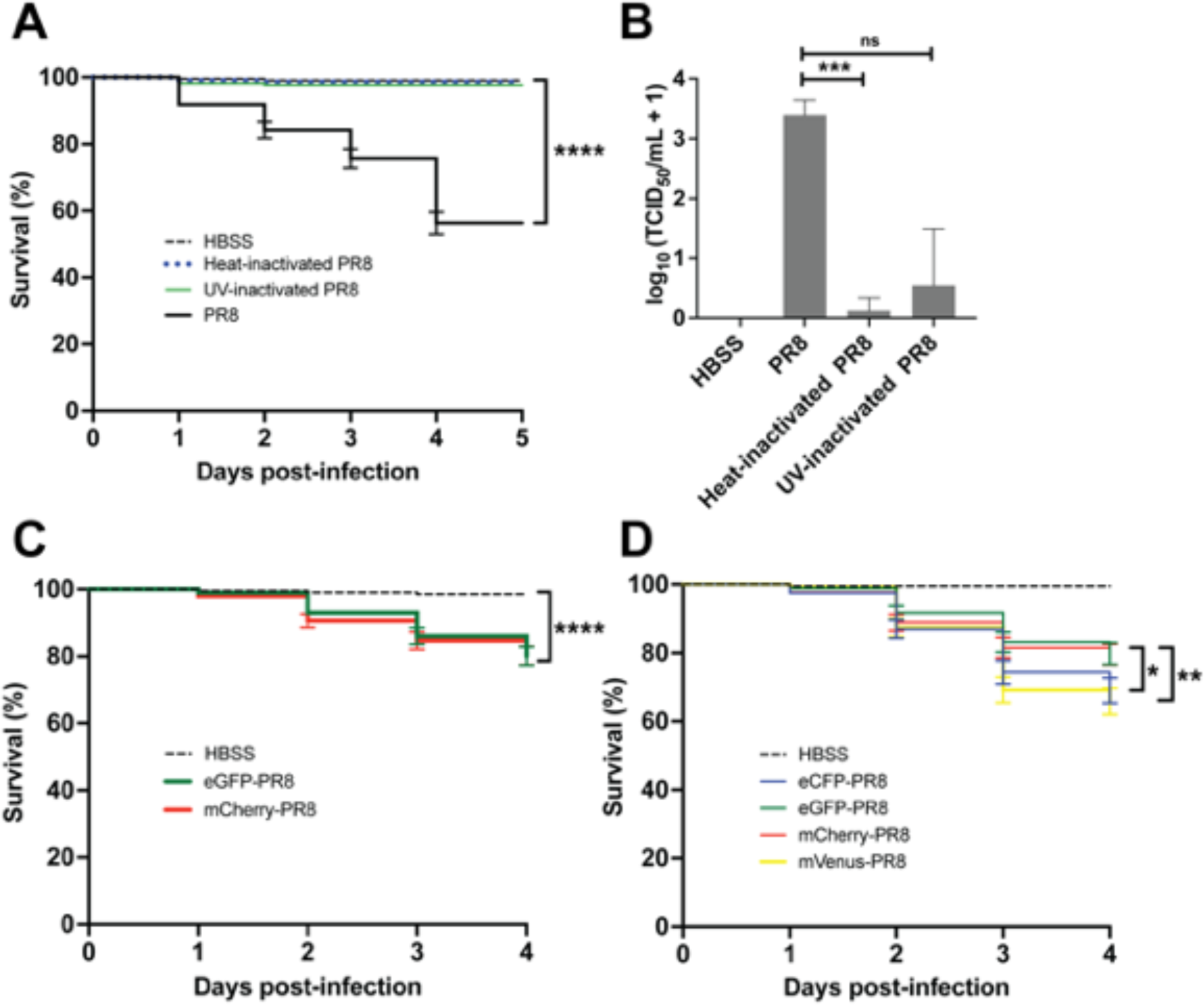
A) Survival of AB larvae injected with live PR8 (8.7 × 102 EID50), heat-inactivated PR8, UV-inactivated PR8 or HBSS (vehicle control) at 2 dpf. Live PR8 infection resulted in significantly lower survival than HBSS, heat-inactivated PR8, or UV-inactivated PR8 (p < 0.0001). B) Viral burden of larvae at 24 hpi with live PR8 (8.7 × 102 EID50), heat-inactivated PR8, and UV-inactivated PR8 at 2 dpf. 24 hpi. Viral burden was different between live PR8 and heat-inactivated PR8 (p = 0.0001). C) Survival of larvae was reduced with eGFP-PR8 and mCherry-PR8 infection compared to HBSS controls (p < 0.0001). D) Survival of Color-flu infected larvae was lower with eCFP-PR8 (p = 0.0288) and mVenus-PR8 (p = 0.0063) infected larvae than eGFP-PR8.

**Supplemental Figure S2:**
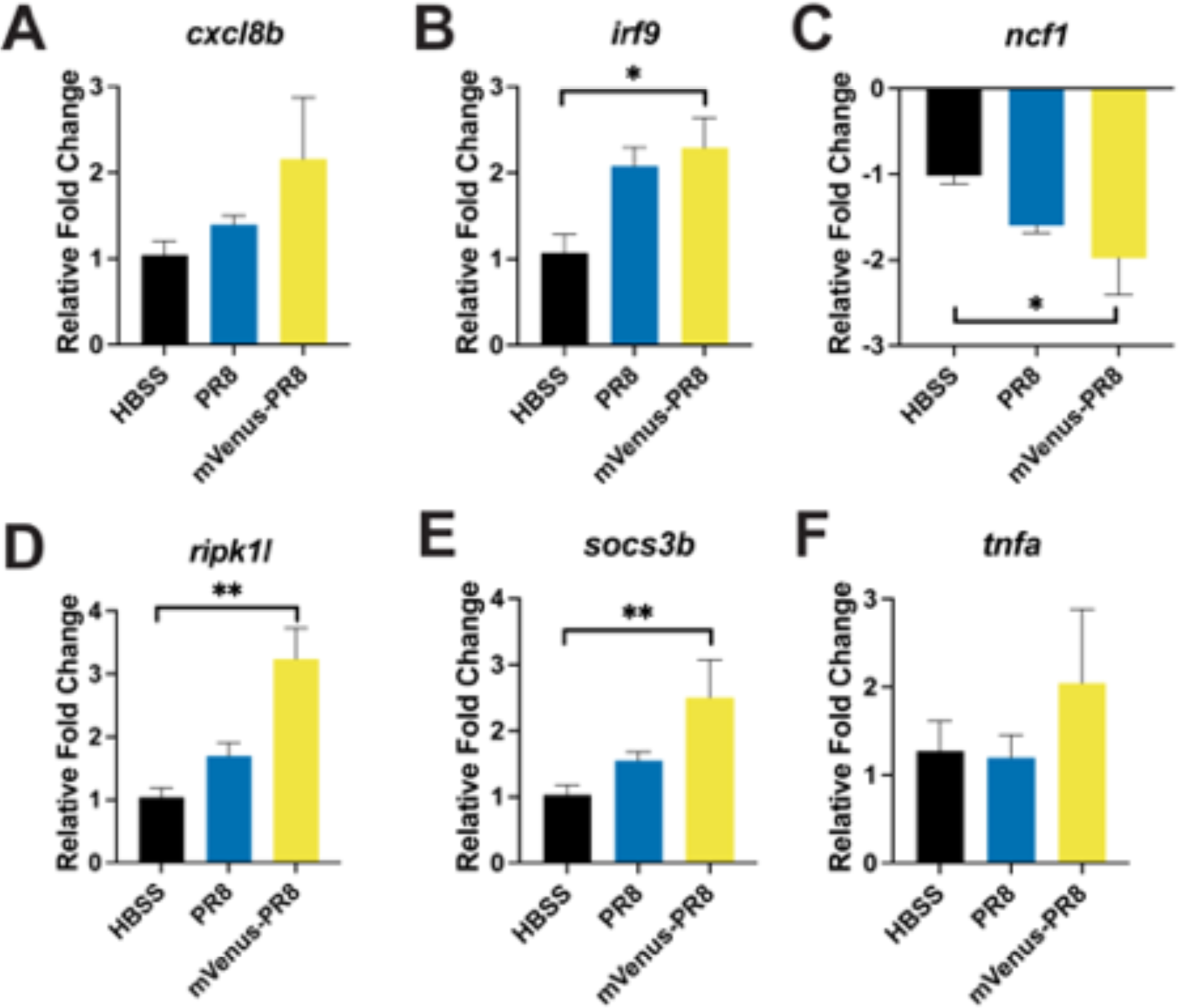
Expression of candidate genes at 24 hpi. Relative fold change of six genes using qRT-PCR (*n* = 4): A) *cxcl8b*; B) *irf9* had increased expression with mVenus-PR8 infection (adjusted p-value = 0.0281); (C) *ncf1* had decreased expression with mVenus-PR8 infection (adjusted p-value = 0.0285); (D) *ripk1l* had increased expression with mVenus-PR8 infection (adjusted p-value = 0.0065); (E) *socs3b* had increased expression with mVenus-PR8 infection (adjusted p-value = 0.0088); and (F) *tnfa*.

**Figure S3:**
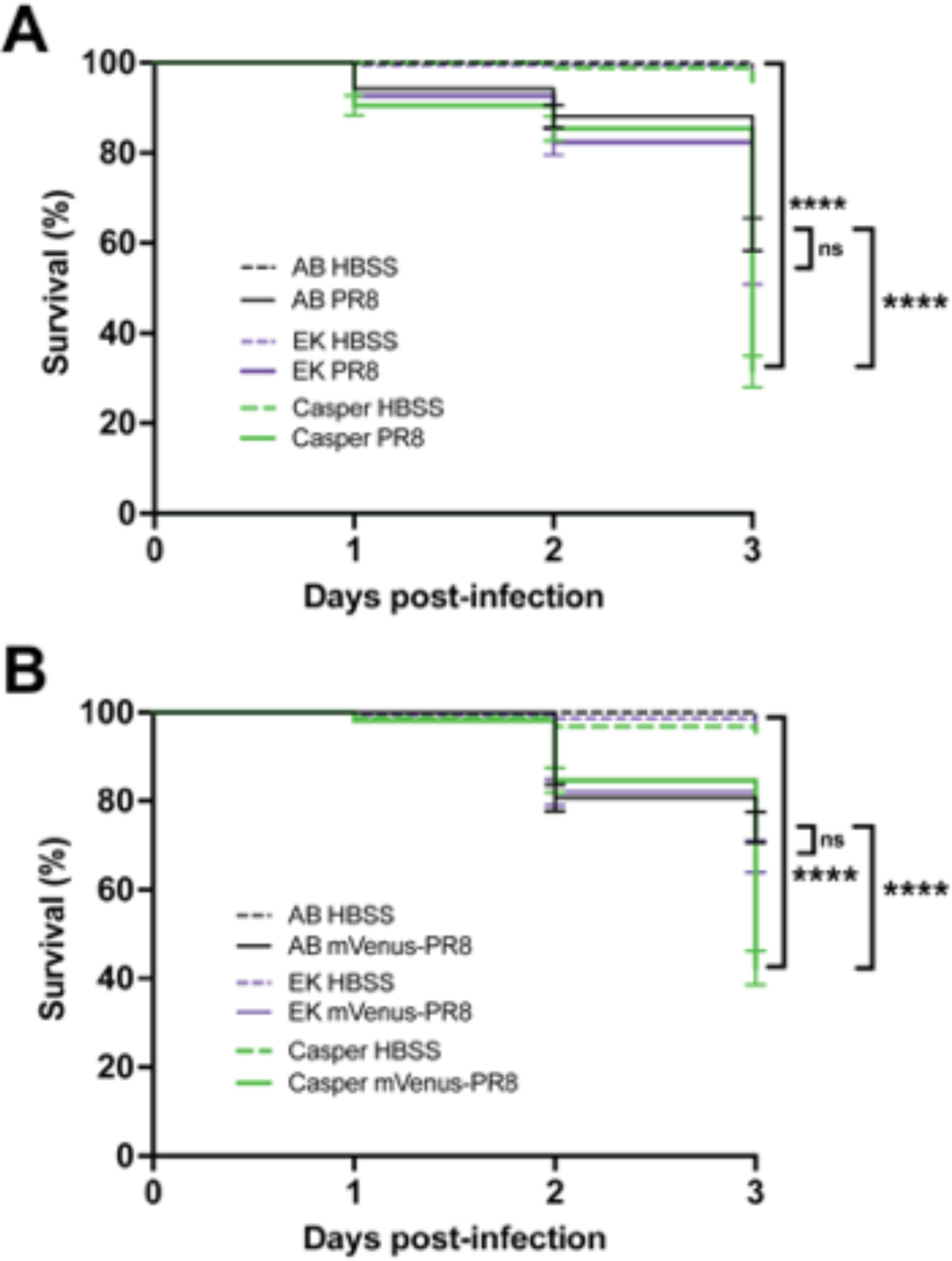
A) Survival of AB, EK and *casper* larvae infected with PR8 in the swimbladder at 4 dpf. Survival was lower in *casper* larvae infected with PR8 than HBSS controls (p < 0.0001). Survival of PR8-infected *casper* lavae was also lower than AB PR8-infected larvae (p < 0.0001). B) Survival of AB, EK and *casper* larvae infected with mVenus-PR8 in the swimbladder at 4 dpf. Survival was lower in *casper* larvae infected with mVenus-PR8 than HBSS controls (p < 0.0001). Survival of mVenus-PR8 -infected *casper* lavae was also lower than AB PR8-infected larvae (p < 0.0001).

**Figure S4:**
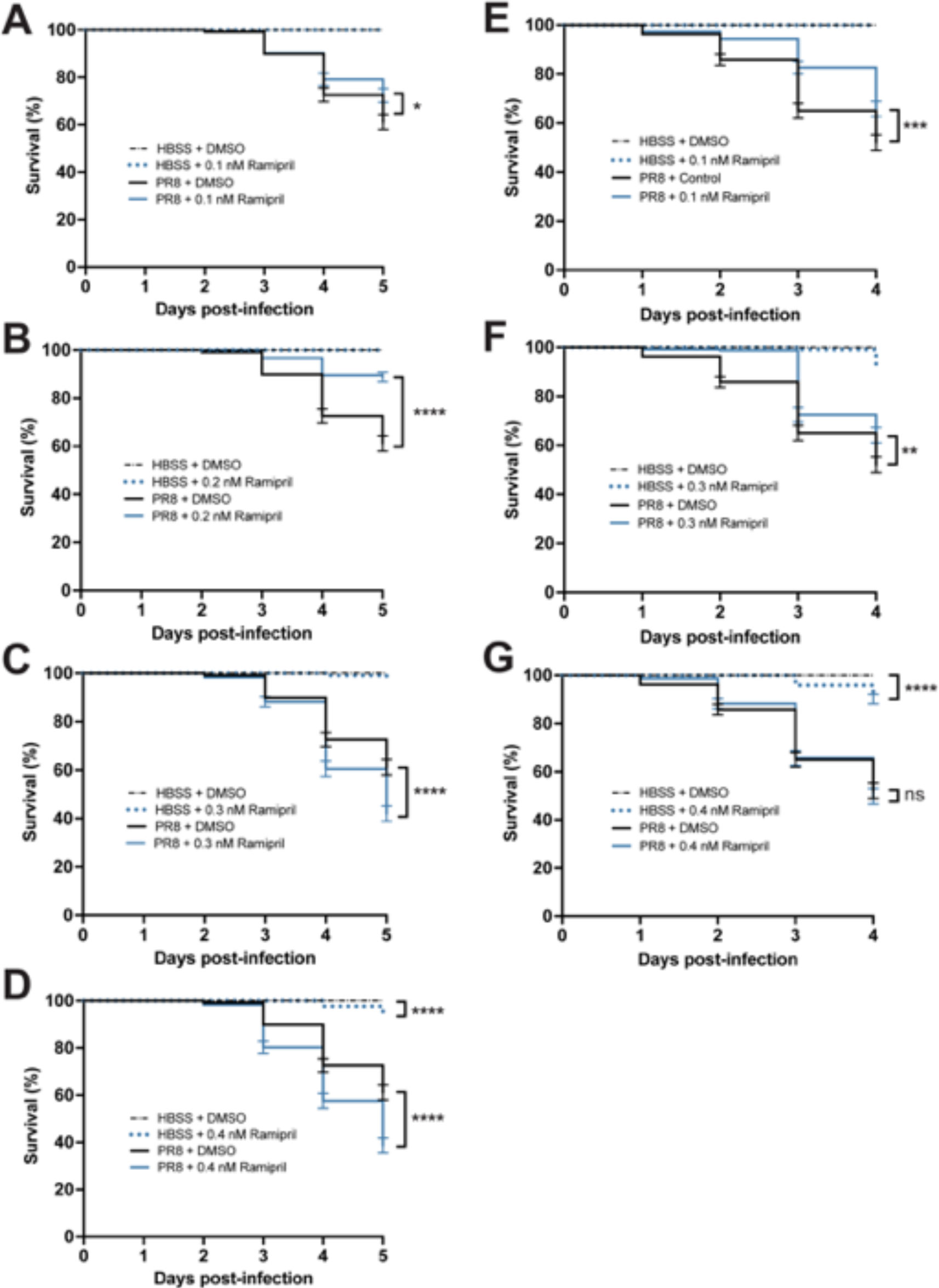
Survival of AB larvae infected with PR8 and treated with DMSO (control) or ramipril. A-D) Survival of larvae infected at 2 dpf and the treated with 0.1, 0.2, 0.3 or 0.4 nM MDIVI-1. Survival was higher in PR8-infected larvae with ramipril exposure at 0.1 nM (p = 0.0157), 0.2 nM, 0.3 nM and 0.4 nM (p < 0.0001) than DMSO controls. Survival was lowered with 0.4 nM ramipril exposure in HBSS controls (p < 0.0001). E-G)) Survival of larvae infected at 3 dpf and MDVI-1 exposure at 0.1 nM (p = 0.0005), 0.3 nM (p = 0.0017). Survival was lowered with 0.4 nM ramipril exposure in HBSS controls (p < 0.0001).

**Figure S5:**
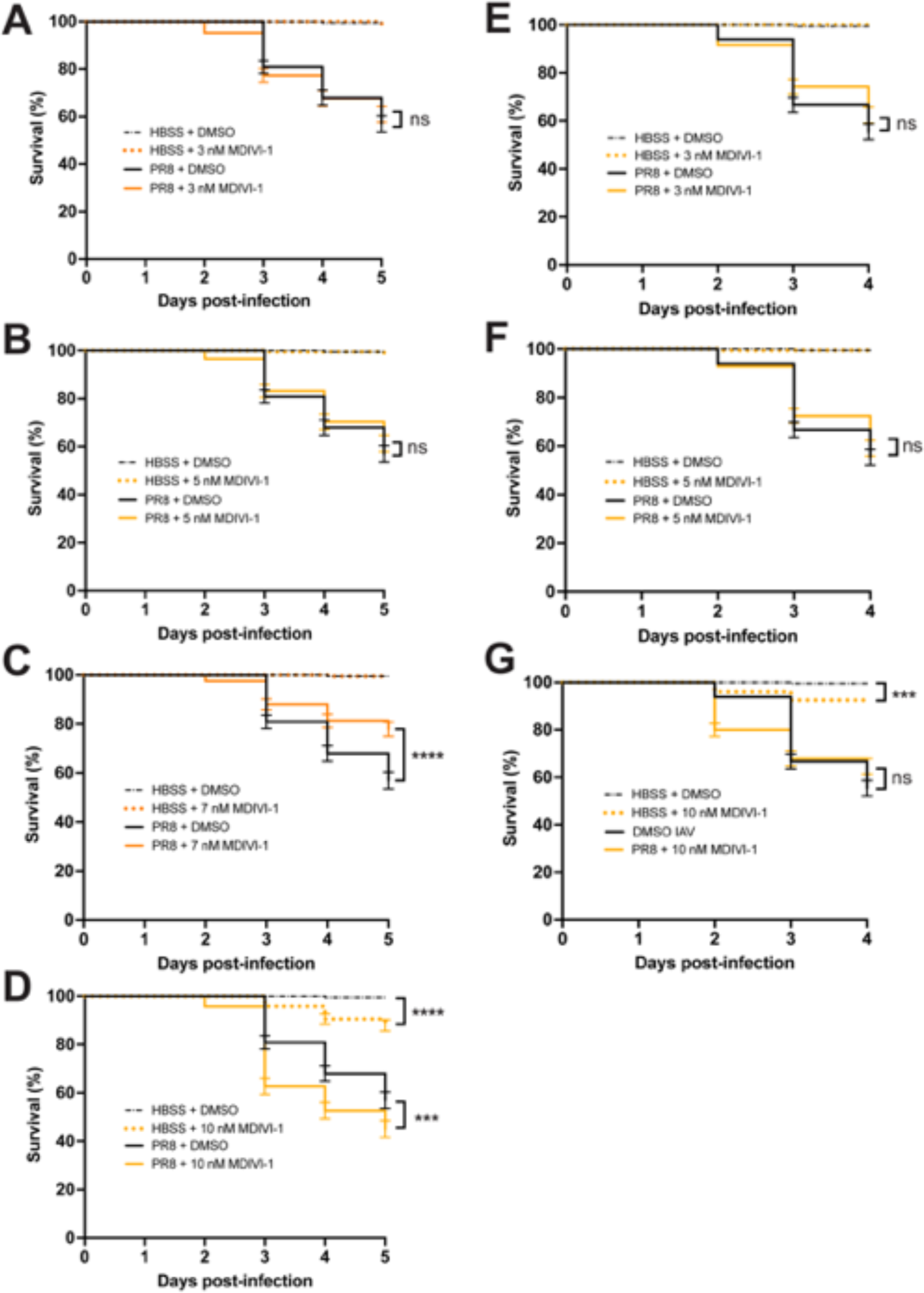
Survival of AB larvae infected with PR8 and treated with DMSO (control) or MDIVI-1. A-D) Survival of larvae infected at 2 dpf and the treated with 3, 5, 7 or 10 nM MDIVI-1. Survival was higher in PR8-infected larvae with 7 nM MDIVI-1 than DMSO controls (p < 0.0001). Survival was lowered with 10 nM MDIVI-1 exposure in HBSS controls (p < 0.0001) and PR8-infected larvae (p = 0.0004). E-G)) Survival of larvae infected at 3 dpf and the treated with 3, 5, or 10 nM MDIVI-1. Survival was lowered with 10 nM MDIVI-1 exposure in HBSS controls (p = 0.0003).

**Supplemental Figure S6:**
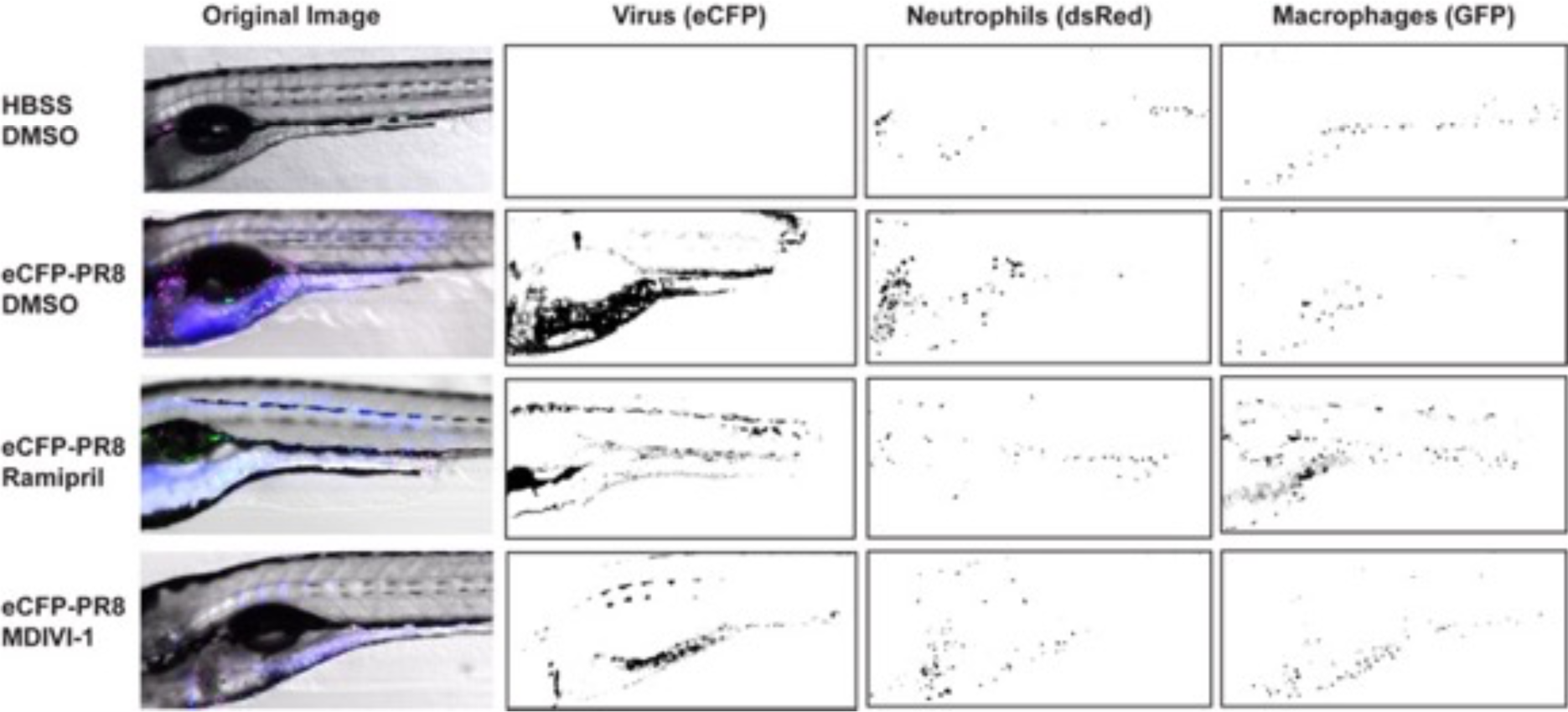
Quantification of virus, neutrophils and macrophages by fluorescent confocal imaging of Tg(*mpeg1:eGFP;lyz:dsRed*) larvae at 48 hours post injection by eCFPPR8 or HBSS following treatment by DMSO, ramipril and MDIVI-1. Representative images and masks used to quantitate the level of eCFP-PR8 virus by pixel intensities in the cyan (emission at 476 nm) channel, number of neutrophils estimated from pixel intensities in the red channel (emission at 583 nm), and number of macrophages estimated from pixel intensities in the green (emission at 510 nm) channel.

